# Prenatal growth map of the mouse knee joint by mean of deformable registration technique

**DOI:** 10.1101/321828

**Authors:** Mario Giorgi, Vivien Sotiriou, Niccolo’ Fanchini, Simone Conigliaro, Cristina Bignardi, Niamh C. Nowlan, Enrico Dall’Ara

## Abstract

Joint morphogenesis is the process during which distinct and functional joint shapes emerge during pre- and post-natal joint development. In this study, a repeatable semi-automatic protocol capable of providing a 3D realistic developmental map of the prenatal mouse knee joint was designed by combining Optical Projection Tomography imaging (OPT) and a deformable registration algorithm (Sheffield Image Registration toolkit, ShIRT). Eleven left limbs of healthy murine embryos were scanned with OPT (voxel size: 14.63¼m) at two different stages of development: Theiler stage (TS) 23 (approximately 14.5 embryonic days) and 24 (approximately 15.5 embryonic days). One TS23 limb was used to evaluate the precision of the displacement predictions for this specific case. The remaining limbs were then used to estimate Developmental Tibia and Femur Maps. Acceptable uncertainties of the displacement predictions were found for both epiphyses (between 0.7 and 1.4 μm, along all directions and anatomical sites) for nodal spacing of 1 voxel. The protocol was found to be reproducible with maximum Modified Housdorff Distance differences equal to 1.9 μm and 1.5 μm for the tibial and femoral epiphyses respectively. The effect of the initial shape of the rudiment affected the developmental maps by 21.7 μm and 21.9 μm for the tibial and femoral epiphyses respectively, which correspond to 1.4 and 1.5 times the voxel size. To conclude, this study proposes a repeatable semi-automatic protocol capable of providing mean 3D realistic developmental map of a developing rudiment allowing researchers to study how growth and adaptation are directed by biological and mechanobiological factors.

## Introduction

Growth and morphogenesis are two fundamental processes which every living system undergo during both the prenatal and juvenile phase. If growth is more related to an increase in size and mass of an organism over a period of time, morphogenesis is the biological process responsible for any organism to develop its shape. Joint morphogenesis is the key process through which the two opposing cartilaginous rudiments of a joint develop their reciprocal and fully functional shapes, and which starts during prenatal joint development. This process is described in details by Pacifici et al., [1] and subsequently updated by Nowlan and Sharpe [2]. The consequences of incomplete or abnormal joint morphogenesis can be very debilitating and may lead to musculoskeletal diseases such as osteoarthritis (OA) [3, 4]. The formation of a skeletal joint is a highly regulated process which is controlled by several biochemical factors (e.g. growth factors, Hox genes) [5]. Moreover, several studies have experimentally found a relationship between prenatal joint motion and physiological joint morphogenesis in mice [6, 7] and chicks [8–11], demonstrating the importance of mechanical loads in the morphogenic process. For example, 2D histological assessment of chick embryos immobilised with neuromuscular blocking agents showed a reduction in width of the intercondylar fossa of the distal femurla and of the proximal epiphysis of the tibiotarsus and fibula during knee joint morphogenesis [10], and up to 50 % reduction in the epiphyseal width of the proximal and distal regions of the knee, tibiotarsus and metatarsus [9]. Despite the clinical relevance of morphogenesis, there is very little understanding about the factors driving this complex process [1]. Mechanobiological growth models have also been used to deepen our understanding on morphogenesis by exploring the role of motion or loading on joint shape [5, 12–15]. However, despite their undeniable importance these models have a series of limitations. For example, the 3D prenatal joint kinematics and kinetics are not measured accurately, and so far only generic loading conditions have been used for the Finite Element (FE) models. Moreover, idealised joint shapes were used instead of realistic shapes. Finally, due to a lack of information, the cascade of biochemical factors determining the amount of biological growth on which the mechanical stimuli operates were extremely simplified and considered to be proportional to the chondrocytes density in the region of interest [12, 13].

This study focuses on defining a methodology for better assessment of realistic shapes in the prenatal joint morphogenesis. Principal Component Analyses (PCA) [16, 17] is the gold standard method for the assessment of shape changes from medical images. However, this method is based on predefined modes of deformation and, due to the lack of large databases of prenatal rudiment shape changes, this approach could not be used for this application. The approach used in this study is based on deformable image registration [18, 19], which, due to the large number of degrees of freedom of the transformation, can feasibly be used to measure the heterogeneous variations of the developing rudiments.

In this study, we developed a repeatable semi-automatic protocol capable of providing a 3D realistic developmental map of the developing mouse knee joint by applying a deformable registration algorithm (Sheffield Image Registration toolkit, ShIRT [18–20]) to 3D images acquired ex vivo using Optical Projection Tomography (OPT) [21]. The robustness and repeatability of the protocol was evaluated with inter- and intra-operator tests. The developed protocol can be used to study the effect of mechanical and biological stimuli on the joint growth and morphogenesis, and to populate and validate computational models for prediction of joint development.

## Materials and methods

### Specimen preparation and imaging with Optical Projection Tomography

The limbs of mouse embryos were stained for cartilage using Alcian Blue (as reported in [22]) and scanned in 3D with Optical Projection Tomography (OPT) [21]. All samples were healthy embryos, either wildtype or heterozygous, for the Pax3 mutation from the Spd (Splotch delayed) strain. Eleven limbs in total were used for this study, six of which were staged as Theiler Stage (TS23 [23] equivalent to approximately 14.5 embryonic days, and the remaining five which were staged as TS24 equivalent to approximately 15.5 embryonic days). One of the six TS23 embryos was scanned twice and the obtained images were registered to evaluate the precision of the deformable registration algorithm used to measure the developmental map [20]. All procedures performed complied with the ethical European Legislation. The project license used was approved by the Home Office and by the Governance Board for Animal Research at Imperial College London.

### Elastic registration protocol

After image reconstruction of each scanned rudiment (NRecon, Bruker microCT, Belgium) the eleven distal femoral and the eleven proximal tibial epiphyses were cropped from the images of the hindlimbs by applying a single level threshold followed by a manual refinement of the initial segmentation based on visual checks in each orthogonal plane (AMIRA software 2017, Thermo Fisher Scientific). The two developmental stages used in this study (TS23 and TS24) were selected due to the fact that at earlier stages (<TS23) incomplete separation of the rudiments did not allow their proper identification, while at later stages (> T24), epiphyseal segmentation was compromised by advanced ossification in the diaphysis. Next, two of the TS23 images (a femoral epiphysis and a tibial epiphysis) were removed from the analysis and used to evaluate the precision of the deformable registration algorithm as described later in this section. A bounding box was then used to crop every remaining specimen by including a similar portion of the diaphysis (Fig 1, A). The analyses were performed on the distal femur and on the proximal tibia to avoid the diaphysis, a region with limited or absent contrast in the OPT images due to the advanced ossification for the TS24 samples. In addition, one TS23 femoral epiphysis was excluded due to incomplete separation of the rudiments. All remaining TS23 tibial (N = 5) and femoral (N = 4) epiphyses were rigidly registered to each TS24 epiphyses through an automatic alignment of the centres of mass (AMIRA software function) followed by a manual adjustment of the orientation to align the main features of the rudiments (see Fig 1, A-B). A total of 25 and 20 rigid registrations were performed for the proximal tibia and the distal femur, respectively. The same bounding box was then used to resample all the images with a Lanczos interpolator [24]. To minimize the imperfections due to manual segmentation, a 3D erosion algorithm of 1 voxel was applied to all the images. Each pair of rigidly registered TS23-TS24 images was then registered with a deformable registration algorithm (ShIRT) [18, 20] in order to compute the displacements at the nodes of an isotropic grid superimposed to the images with nodal spacing (NS) equal to one voxel (14.63μm), for maximizing the number of degrees of freedom in the registration displacement map. In order to analyse only the results obtained for the rudiments, every cell of the grid with all nodes outside the TS23 binary image was removed by using a custom-made script (Matlab, The MathWorks, Inc.) [25].

**Fig 1.**
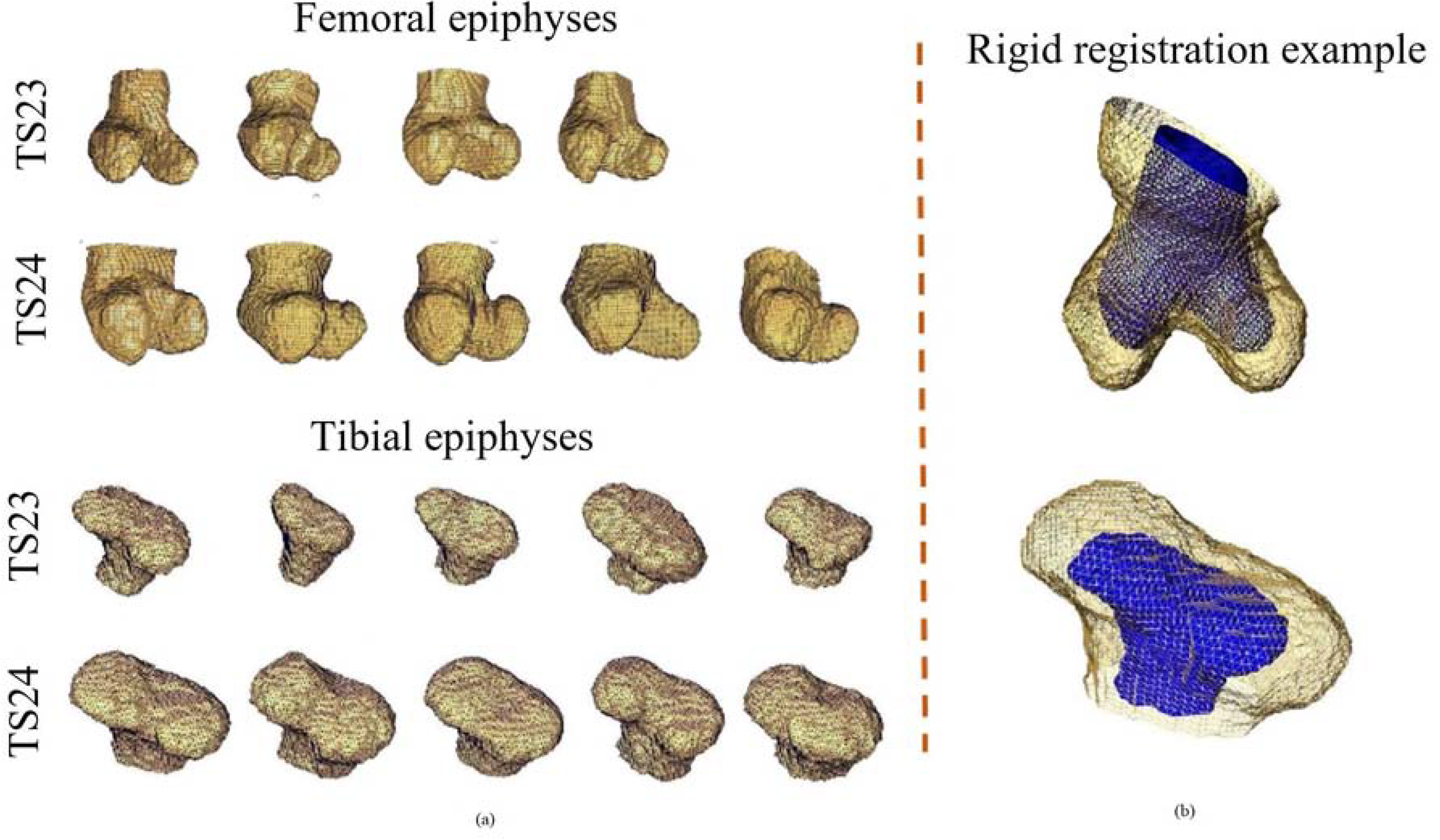
Reconstructed epiphyses used for the study and example of rigid registration. (a) femoral and tibial epiphyses (TS23 and TS24) used for the study; (b) example of rigidly registered femoral (top) and tibial (bottom) epiphyses. The TS23 epiphyses are represented in blue and the TS24 epiphyses in yellow.

The excluded TS23 sample was used to evaluate the uncertainties in the displacement predictions by registering two pairs of repeated scans of the whole prenatal femur and tibia following a procedure used for different bone structures [26, 27]. The precision of the method was evaluated for the three Cartesian directions using the standard deviation of the displacement components over the whole registration grid for tibia and femur. Uncertainties equal to 1.4 μm, 1.4 μm and 1.3 μm were found for the tibia along X, Y and Z directions, and uncertainties of 1.0 μm, 1.0 μm and 0.7 μm were found for the femur along X, Y and Z directions, respectively. These errors were considered acceptable for this application where displacements were at least one order of magnitude larger than the measured uncertainties.

Considering the absence of an in vivo longitudinal imaging modality for OPT measurements and the intrinsic variability of the shape of the rudiments at TS23 and TS24 (Fig 1, A), the effect of including different input images on the final developmental map was estimated as follows (overview of the procedure in Fig 2).

**Fig 2.**
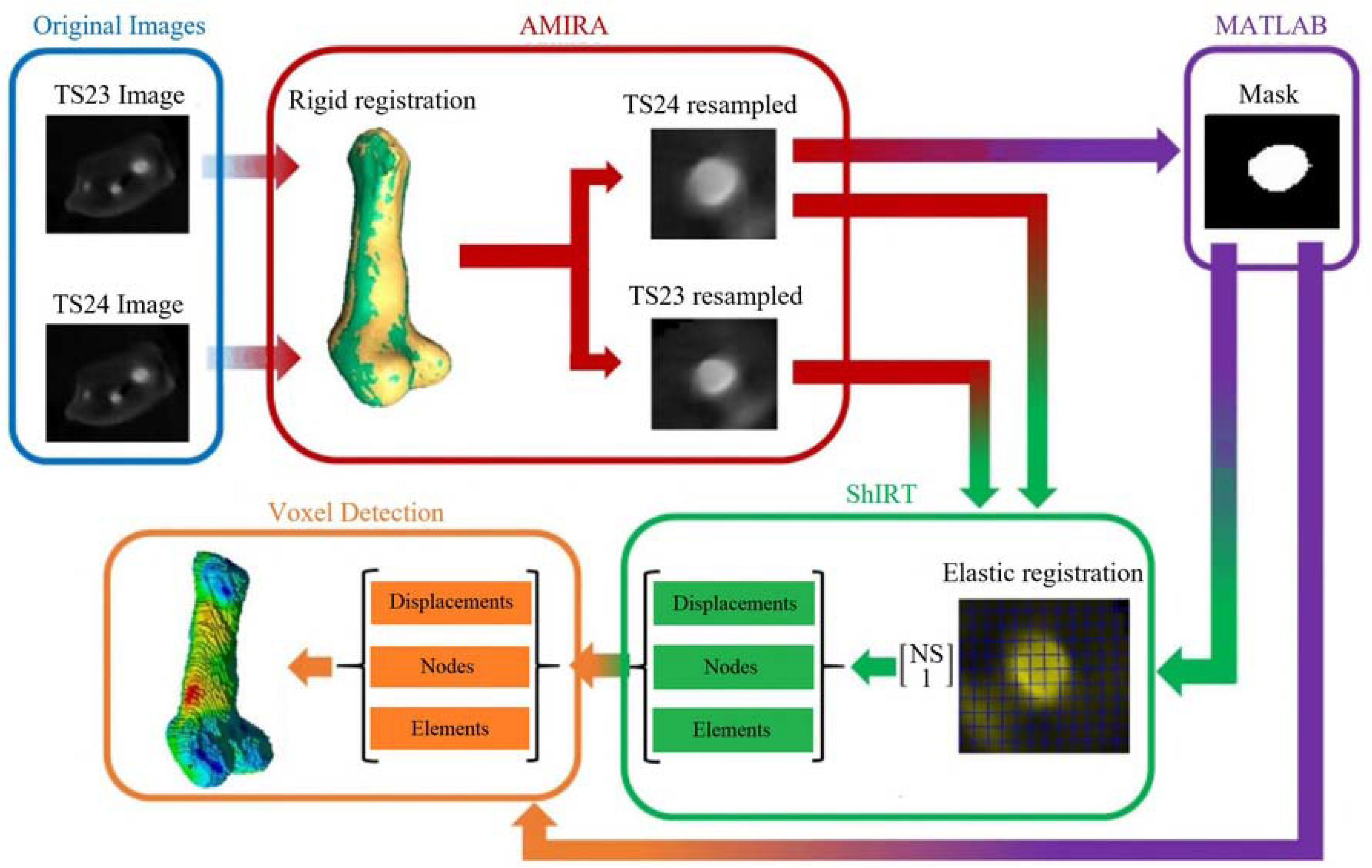
Methodological pipeline. Two OPT images at different developmental stages were acquired and rigidly registered. The new resampled images, together with a B/W image of the latest developmental stage (TS24) were then given as input to ShIRT. The calculated displacement were then filtered thought the Voxel detection toolkit.

Twenty displacement maps were generated by deformable registration of four randomly picked stack of images from the TS23 tibial epiphysis group with every image of the TS24 tibial epiphysis group. The obtained displacement values were averaged in order to generate a mean growth map (from now on referred to as “Developmental Tibia Map”, DTM). Furthermore, the average of five displacement maps obtained by registering the remaining image of the TS23 tibial epiphyses group with each one of the images in the TS24 tibial epiphysis group was computed in order to evaluate a control map (from now on referred to as “Single Tibia Map”, STM) (Fig 3, A). The same procedure was then applied to the femur samples in order to generate the “Developmental Femur Map” (DFM), and “Single Femur Map” (SFM) (see Fig 3, B).

**Fig 3.**
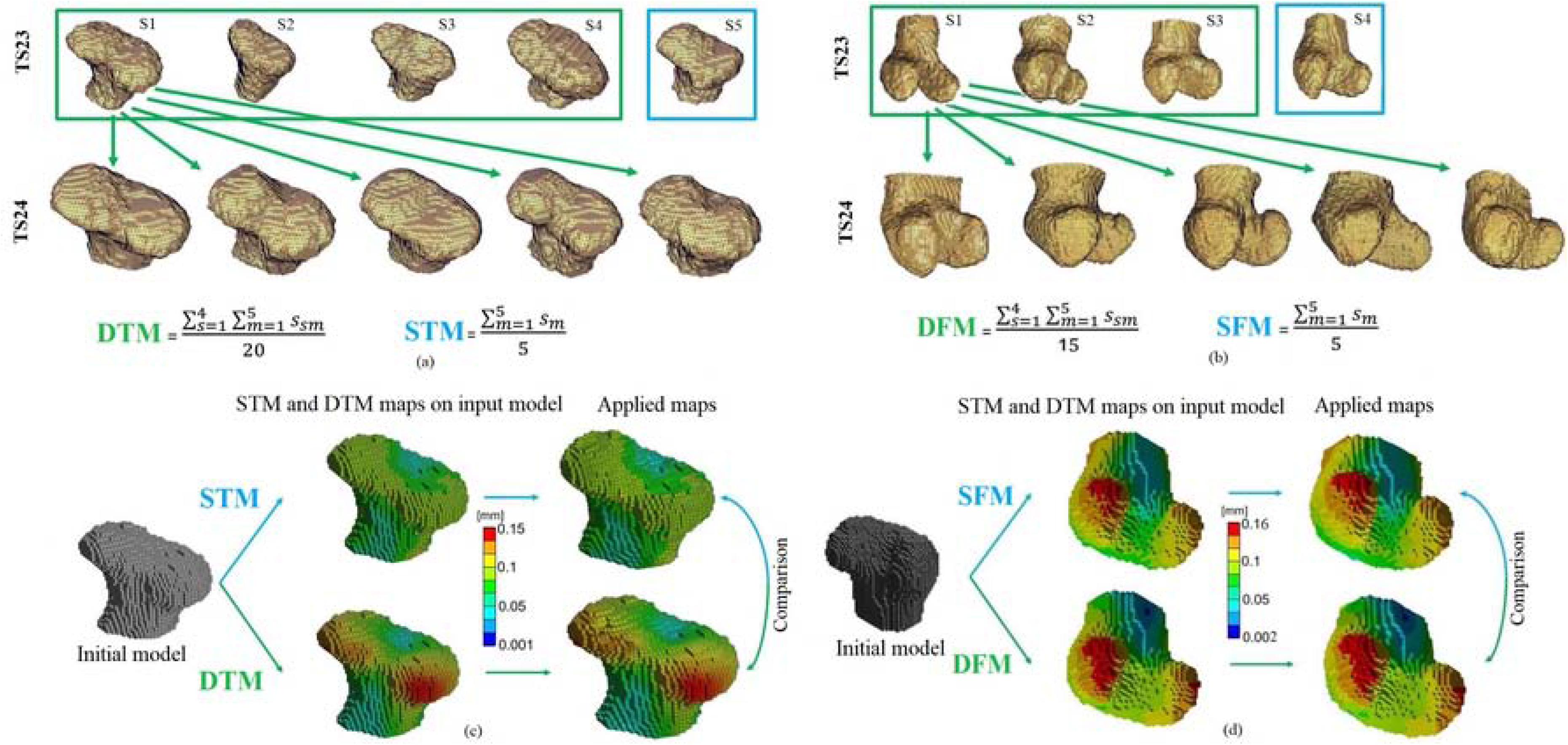
Schematic representation for the generation of both tibial and femoral maps. (a) Schematic representation of the twenty displacement maps (green box) and five displacement maps (blue box) generated by deformable registration for the tibia. The obtained maps were then averaged in order to generate a Developmental Tibia Map (DTM), and a Single Tibia Map (STM); (b) Schematic representation of the fifteen displacement maps (green box) and five displacement maps (blue box) generated by deformable registration for the femur. The obtained maps were then averaged in order to generate a Developmental Femur Map (DFM), and a Single Femur Map (SFM); (c) qualitative comparison between the DTM and the STM showing similar developmental patterns; (d) qualitative comparison between the DFM and the SFM showing similar developmental patterns.

For the visualization of the displacement maps for tibia and femur, the registration grids of the TS23 control specimens were converted into meshes of 8-node hexahedron elements, and the “Developmental Maps” and “Single Maps” were applied as kinematic boundary conditions. A finite element (FE) software package (ANSYS, Mechanical APDL v.15.0, Ansys Inc, USA) was used to visualize and compare the maps.

## Analyses and comparison of the growth maps

Two different analyses, briefly described below, were performed in this study: 1) Comparison of the Developmental and Single Maps for tibia (DTM, STM) and femur (DFM, SFM); 2) Evaluation of the protocol repeatability.

### Comparison of the Developmental and Single Maps for both anatomical sites

For both epiphyses, the displacement distributions of the Developmental Map and the Single Map were qualitatively compared after their application to the TS23 control FE mesh. Then, all the surface nodes of the new rudiment shapes were extracted using a custom-made script (Matlab, The MathWorks, Inc.), and their differences quantified using the Modified Hausdorff Distance (MHD) *[28]* and the Average Displacement Distance (ADD). For visualization purposes, two STL surfaces were then generated by using the ball-pivoting technique [29] available in Meshlab.

### Evaluation of the protocol repeatability

The repeatability of the protocol was evaluated through intra- (three repetitions) and inter-operator (one expert operator and two operators who followed guidelines) tests. For the tibial epiphyses, both tests were performed, while only the intra-operator test was performed on the femoral epiphyses. The part of the protocol mostly affected by operator decisions is the initial manual orientation of the rudiments. Therefore, the protocol was repeated from the initial upload of the raw images into Amira until the generation of the DTM, STM, DFM and SFM. The standard deviation of the MHD and ADD values for the three intra-operator repetitions and for the three inter-operator repetitions were computed.

## Results

### Comparison of the Developmental and Single Maps for both anatomical sites

A qualitative comparison between the Developmental Tibia and Femur Maps (DTM, DFM), and between the Single Tibia and Femur maps (STM, SFM) showed similar developmental patterns, with higher displacement values on the lateral and medial condyles of both rudiments compared to the inter-condylar region (Fig 3, CD). On a more quantitative aspect, displacements up to 150 μm and 100 μm were observed in the condyles region for the Developmental Tibia and Single Tibia maps respectively (DTM, STM), and displacements of approximately 60 μm were measured in the intercondylar region for both maps (Fig 3, C). Similarly, displacements of up to 160 μm were observed in the condyles region for the Developmental Femur and Single Femur Maps (DFM, SFM) respectively, and displacements of approximately 80 μm were measured in the inter-condylar region for both maps (Fig 3, D). When the effect of the initial shape of the rudiment over the developmental maps was quantified, MHD value of 21.7 μm and ADD values of 32.5 μm along X axis, of 35.6 μm along Y axis, and of 29.2 μm along Z axis were found between the Developmental Tibia Map (DTM) and the Single Tibia Map (STM) (Table 1).

**Table 1.** Modified Hausdorff Distance (MHD) and Average Displacement Distance (ADD) values for all performed analyses for both tibial and femoral epiphyses. Values expressed in μm.

| Tibial Epiphysis | MHD | MHD (range) | MHD (max difference) | ADD | ADD-X | ADD-Y | ADD-Z |
| --- | --- | --- | --- | --- | --- | --- | --- |
| Inter-Operator Test | $20.0 \pm 0.2$ | 19.8 – 20.2 | 0.4 | $28.8 \pm 0.2$ | $28.6 \pm 0.5$ | $31.9 \pm 0.6$ | $25.8 \pm 0.3$ |
| Intra-Operator Test | $21.1 \pm 0.3$ | 22.2 – 22.4 | 0.4 | $29.9 \pm 0.8$ | $30.0 \pm 2.2$ | $33.7 \pm 1.6$ | $28.5 \pm 1.0$ |
| Femoral Epiphysis | MHD | MHD (range) | MHD (max difference) | ADD | ADD-X | ADD-Y | ADD-Z |
| Intra-Operator Test | $22.8 \pm 0.7$ | 22.3 – 23.4 | 1.1 | $34.2 \pm 0.4$ | $34.8 \pm 0.8$ | $34.8 \pm 0.1$ | $31.9 \pm 1.4$ |

When the same analysis was performed on the femoral epiphyses an MHD of 21.9 μm and an ADD of 34.2 μm, 34.9 μm, and 30.3 μm were found along X, Y, and Z axes respectively (Table 1). A qualitative comparison between the Developmental Tibia Map (DTM) and the Single Tibia Map (STM), and between Developmental Femur Map (DFM) and the Single Femur Map (SFM) is reported in Fig 4, A.

**Fig 4.**
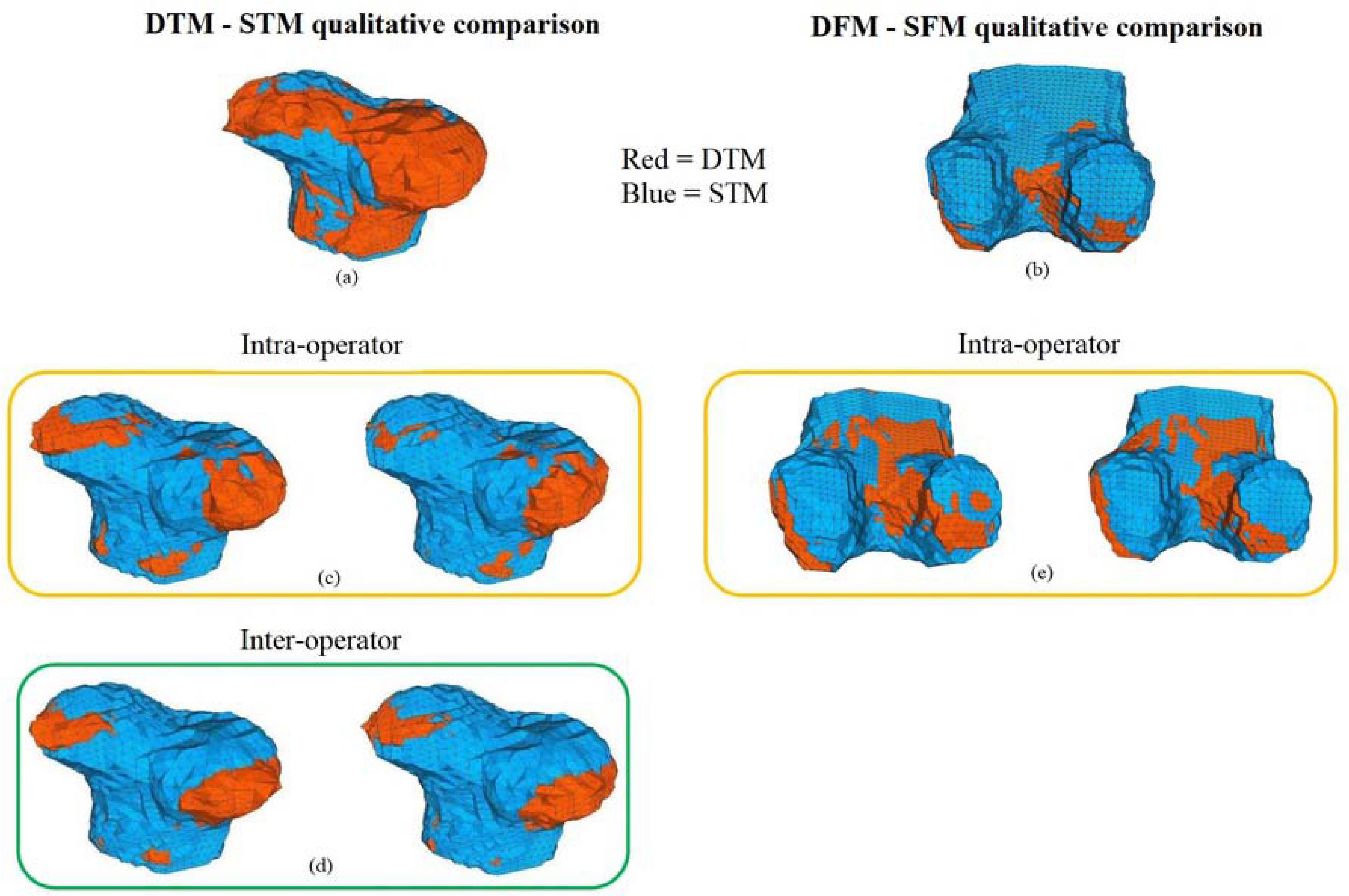
Qualitative comparison between maps. (a) Qualitative comparison between DTM (red) and STM (blue). (b) Qualitative comparison between DFM (red) and SFM (blue). (c) Qualitative comparison between the DTM (red) and STM (blue) for the two tibial femoral intra-operator tests. (d) Qualitative comparison between the DTM (red) and STM (blue) for the two tibial femoral inter-operator tests. (e) Qualitative comparison between the DFM (red) and SFM (blue) for the two tibial femoral intra-operator tests.

### Evaluation of the protocol repeatability

For the tibial epiphyses, MHD values of 22.1±0.4 μm and 20.0±0.3 μm were measured for the intra- and inter-operator test, respectively (see Table 1). A similar MHD intra-operator value was measured for the femoral rudiment (22.5±0.8 μm) (see Table 1). For intra-operator assessment of the tibial rudiment, ADD values of 30.0±2.2 μm, 33.7±1.6 μm, and 28.5±1.1 μm along X, Y, and Z axis respectively were calculated (see Table 1). Similar values were observed for the inter-operator test performed on the same rudiment with 28.6±0.6 μm, 31.9±0.6 μm, and 25.8±0.4 μm along X, Y, and Z axis, respectively (see Table 1). ADD values for the femoral rudiment, showed intra-operator values of 34.8±0.9 μm, 34.8±0.1 μm, and 31.9±1.4 μm along X, Y, and Z axis respectively (see Table 1). A qualitative comparison between intra- and inter-operator generated maps for both epiphyses is reported in Fig 4, C-E.

## Discussion

The aim of this study was to develop a repeatable semi-automatic protocol capable of providing a 3D realistic developmental map of the tibial and femoral developing rudiments in a prenatal mouse model. This was achieved by combining OPT imaging [21] and a deformable registration algorithm [18–20].

The process of joint morphogenesis is key for physiological skeletal development. However, due to its complexity, there is very little understanding about the factors driving it [1]. Computational models have been used to deepen our understanding on the importance of fetal movement during development [12–14], but, these models are usually based on idealized shapes. The protocol developed in this study can be used to quantify the shape changes of a developing rudiment and provide 3D realistic displacement maps for studying the joint development, at different stages, in healthy or diseased animals and to populate computational models to study morphogenesis and its dependency on mechanical and biological stimuli. The protocol was found to be strongly reproducible for both epiphyses, with small intra- (SD of ADD below 2.2 μm for tibia and 1.4 μm for femur) and inter-operator (SD of ADD below 0.6 μm for tibia) displacement uncertainties (Table 1). The high reproducibility of this protocol enables researchers not familiar with elastic registration to generate reliable developmental maps. In addition, the maximum reproducibility error measured in terms of ADD (2.2 μm) suggests that this method is suitable to study all deformations of at least one order of magnitude higher (all deformations higher than 22 μm) and this method could be used in the future to generate growth maps for earlier stages where the experimentally observed deformations have not yet been quantified precisely. The robustness of the method is also highlighted by the similarity between the reproducibility errors for the two anatomical rudiments and among the Cartesian directions.

For the developmental stages analysed (TS23 and TS24), the generated maps for both anatomical sites showed higher growth corresponding to the condyle regions and lower growth in the intercondylar fossa. In addition, the DTM and the DFM showed mean displacement values of 75 μm and 81 μm, with maxima values of 150 μm and 160 μm for the tibia and femur respectively (Fig 3, C, D). When the DTM and DFM were qualitatively compared with the STM and SFM, the analyses showed, as expected, similar but not identical developmental patterns (Fig 3, C, D). Maximum MHD between the measured displacement maps of 21 μm and 22 μm were found for the tibial and femoral maps, showing the variability of the growth among specimens. This value, ten times higher than the estimated uncertainties, underlines the important role that the input images play. In fact, the high variability in the developmental maps found in this study is probably based on the differences in shape found especially at TS23 for both tibial and femoral rudiments (see Fig 1 for examples). Such variability could be probably reduced by increasing the sample size, to account for intrinsic differences among the specimens, or by extending the OPT imaging to in vivo application, something which is not possible right now.

This study has mainly two limitations. Firstly, the analyses were performed on the distal femur and on the proximal tibia only due to the limited or absent contrast in the OPT images due to the advanced diaphysis ossification for the TS24 samples. A combination between OPT and micro-CT (micro-computer tomography) imaging could help to overcome this problem and allow the application of this protocol to the whole rudiments. Secondly, a more comprehensive developmental map including earlier and later developmental stages could not be generated. The former because at earlier developmental stages the joints were not fully cavitated, making very difficult the identification of specific joint segments. The latter due to the advanced ossification process, which started involving the epiphyses. Finally, the analyses was performed on only 4 or 5 specimens. As the final goal is to create a mean biological growth map, including more specimens (and therefore more registrations) would reduce the influence of the differences in initial shape of the joints at the two TS.

In conclusion, in this study we have shown how a combination of OPT imaging and deformable registration can be used to generate 3D realistic transformations of a developing rudiment in the prenatal mouse knee joint. The method is highly reproducible and will allow us to study how growth and adaptation are directed by biological and mechanobiological factors. Moreover, the realistic shapes can be used to generate more accurate computational models capable of exploring the influence of both physiological and non-physiological mechanical conditions on the process of morphogenesis and joint development.

## Acknowledgements

The authors gratefully acknowledge Dr. Gareth Fletcher for implementing the DVC service, Sara Oliviero and Maria Cristiana Costa for helping with the repeatability tests.

